# An Oculometrics-based Biofeedback System to Impede Fatigue Development during Computer Work: a Proof-of-Concept Study

**DOI:** 10.1101/563932

**Authors:** Ramtin Zargari Marandi, Pascal Madeleine, Øyvind Omland, Nicolas Vuillerme, Afshin Samani

## Abstract

A biofeedback system may objectively identify fatigue and provide an individualized timing plan for micro-breaks. We developed and implemented a biofeedback system based on oculometrics using continuous recordings of eye movements and pupil dilations to moderate fatigue development in its early stages. Twenty healthy young participants (10 males and females) performed a cyclic computer task for 31-35 min over two sessions: 1) self-triggered micro-breaks (manual sessions), and 2) biofeedback-triggered micro-breaks (automatic sessions). The sessions were held with one-week inter-session interval and in a counterbalanced order across participants. Each session involved 180 cycles of the computer task and after each 20 cycles (a segment), the task paused for 5-s to acquire perceived fatigue using Karolinska Sleepiness Scale (KSS). Following the pause, a 25-s micro-break involving seated exercises was carried out whether it was triggered by the biofeedback system if the fatigue state (KSS≥5) was detected in automatic sessions or by the participants in manual sessions. National Aeronautics and Space Administration Task Load Index (NASA-TLX) was administered after sessions. The functioning core of the biofeedback system was based on a Decision Tree Ensemble model for fatigue classification, which was developed using an oculometrics dataset previously collected during the same computer task. The biofeedback system identified fatigue states with a mean accuracy of approx. 70% and remained robust against circadian rhythms. Perceived workload obtained from NASA-TLX was significantly lower in the automatic sessions compared with the manual sessions, *p*=0.01 Cohen’s *d*=0.89. The results give support to the robustness and effectiveness of integrating oculometrics-based biofeedback in time planning of micro-breaks to impede fatigue development during computer work.

## Introduction

Fatigue is often reported by computer users [1,2] and associated with compromised performance that may result in accidents [3] as well as in the development of musculoskeletal and psychological disorders [2,4]. However, fatigue development is sometimes inevitable due to inflexible work regulations and schedules [2]. Two important issues among all should be addressed in work-related fatigue [2]. First, the regular work-rest schedules may ignore inter-individual differences in the manifestation of fatigue patterns [5,6]. Second, fatigue progression in its early stages may not necessarily lead to a significant loss of performance and thus not easily detectable from performance measures [7,8].

Implementing micro-breaks, i.e. short pauses without major interruption, at work is suggested to mitigate fatigue and preserve the performance in a safe level [9,10]. In addition, micro-breaks have been reported to improve mental focus [11]. It is plausible that micro-breaks can reduce discomfort especially during computer work (e.g. [12]), however, the cognitive impacts of micro-breaks requires further investigations [13]. Optimal design of micro-breaks for an individual requires monitoring fatigue status and acquisition of objective information associated with fatigue [14–17]. The objective information should be provided in an unobtrusive manner to avoid any disturbance to work [14–17]. Oculometrics are believed to be an enriched source of cognitive information and can be achieved by the promising technology of eye tracking [18–22]. The oculometrics may represent the underlying neural mechanism in control and regulation of the eye movements during fatigue development [23]. Recent findings show that the development of fatigue may manifest earlier in the oculometrics than in physical and cognitive performance in various tasks including computer work [18] [24]. Thus, oculometrics are promising for early detection of fatigue.

Effective planning of micro-breaks requires appropriate choice of the period, frequency, and the activity during the micro-breaks [25,26]. These parameters are dependent on the tasks and individuals [27]. Specifically, the frequency of micro-breaks may be determined individually based on oculometrics as sensitive metrics to fatigue development. The aim of this study was to develop a biofeedback system based on oculometrics to provide personalized information on when to apply micro-breaks.

A biofeedback system is commonly comprised of an acquisition system to record physiological data from an individual, a processing unit to interpret the data, and an interface, e.g. a computer screen, to deliver information in real-time according to the processed data to the individual. The underlying idea of biofeedback is to provide cognitive interventions to enhance self-awareness to improve health and performance [28,29]. There are different applications for biofeedback [28,29], e.g. decrease of the muscular load during computer work [14,15], adjustment of the mental load of computer games using physiological signals including pupil diameter changes [30], mental training in competitive sports [31], and counteracting stress and anxiety [32–34].

The proposed biofeedback system in this study was designed to alert participants to take a micro-break during computer work based on oculometrics in a statistical model describing fatigue states. Of note, counteracting fatigue within this framework of cognitive intervention is favorable from the consumption of psychoactive drugs, e.g. caffeine in association with health risks [35]. We hypothesized that the oculometrics-based biofeedback system would impede fatigue development without compromising the performance of computer work.

## Methods and materials

### Participants

Twenty participants, 10 females and 10 males, aged 26 (*SD* 3) years old with the height of 1.72 (*SD* .10) m, and the body mass of 69 (*SD* 15) kg were recruited. All participants had normal or corrected-to-normal vision (self-reported and examined by Snellen chart). The participants were familiar with computer work and used their right hand as their dominant side for computer mouse. Participants were asked to abstain from alcohol for 24 h, and caffeine, smoking and drugs for 12 h prior to experimental sessions. The participants reported at least 6 h (mean 7.6 ± 0.8 h) of night sleep before the experimental session. The Fatigue Assessment Scale (FAS) [36] and the Visual Fatigue Scale (VFS) [37] were administered respectively to exclude participants suffering from chronic fatigue and eye strain. No participant was found with chronic fatigue and eye strain. Written informed consent was obtained from each participant. The experiment was approved by The North Denmark Region Committee on Health Research Ethics, project number N-20160023 and conducted in accordance with the Declaration of Helsinki.

### Experimental approach

A counterbalanced-measures design was employed to investigate the effectiveness of biofeedback-triggered micro-breaks in comparison with self-triggered micro-breaks. To do this, two experimental sessions were conducted in two counterbalanced sessions (days) with one-week inter-session interval.

#### Computer task

In both experimental sessions, participants were asked to perform a cyclic computer task [18] for approx. 31-35 min (Fig 1). The task [38] developed on MATLAB R2018a (The MathWorks, Natick, MA) was displayed on a 19-in screen (1280×1024 pixels, refresh rate: 120Hz) located approx. 58 cm in front of a sitting participant subtending 27°×22° of visual angle (Fig 1b). The task involved 180 cycles each taking approx. 10 s (corresponding to methods times measurement (MTM-100) [39,40]). Each cycle began by memorizing a random pattern of connected points with different shapes presented on a computer screen. The order of connecting points was determined by a textual cue displayed under the pattern indicating the starting point. It was followed by a washout period where no pattern was displayed, and the participants were instructed to keep their gaze on a cross in the center of screen. The cycle continued by the presentation of the doubled-size replica of the pattern without connecting lines. To redraw the lines and replicate the presented pattern, participants were required to click on a sequence of the pattern points as targets. Once the allocated time to replicate the pattern passed, a new cycle with a different pattern was presented. In this design, the perceived level of fatigue based on Karolinska Sleepiness Scale (KSS) [41], was indicated by the participants after each 20 cycles, i.e. segment, in five seconds (KSS pause). The KSS can be rated from one (extremely alert) to 10 (extremely sleepy, can’t wait to sleep).

**Fig 1.**
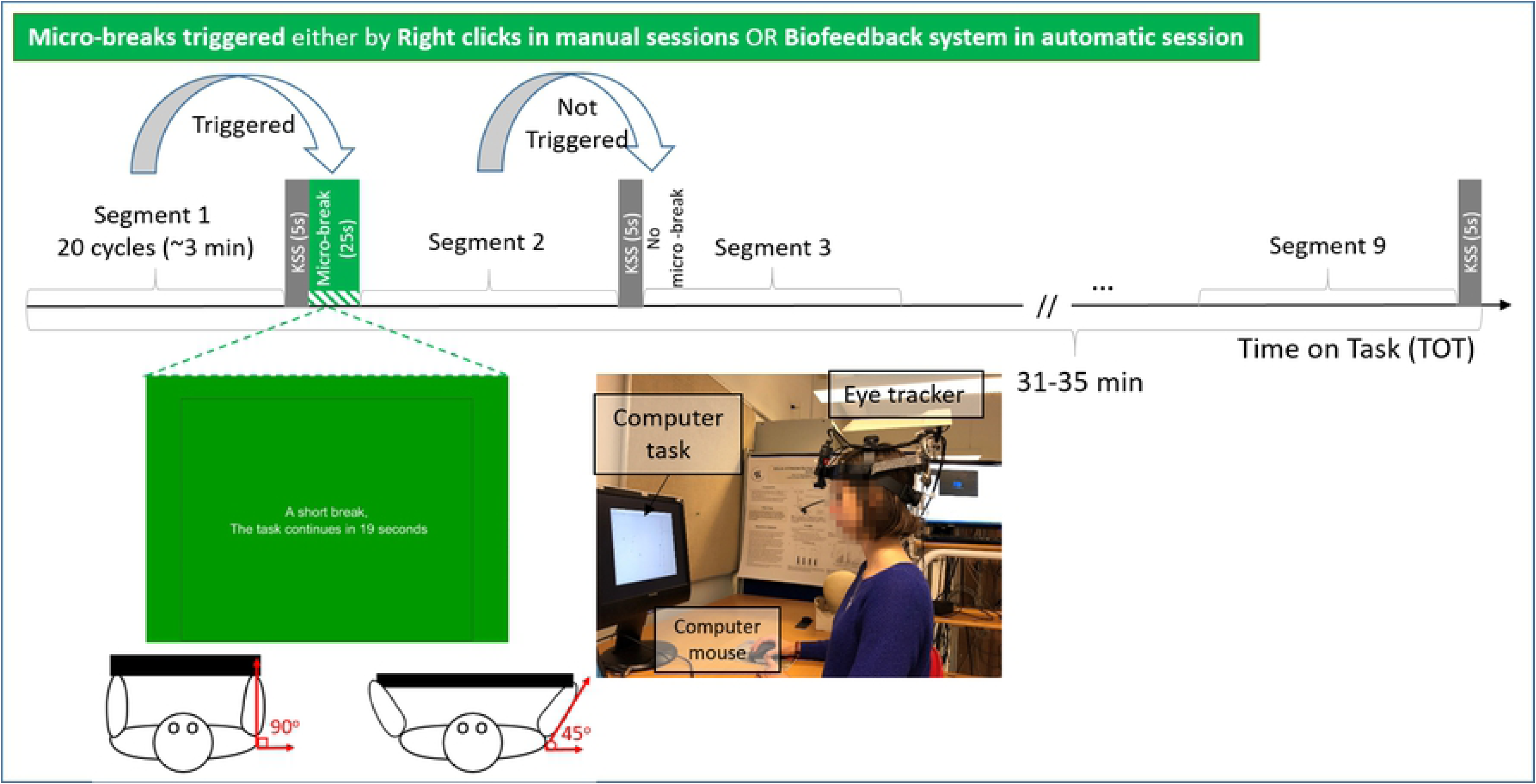
The task timeline in manual and automatic sessions. A schematic view of the exercise and screen information during a micro-break, and the experiment set-up.

#### Micro-breaks

Each experimental session involved either self-triggered or the biofeedback-triggered micro-breaks, respectively termed as manual and automatic sessions. In the manual sessions, the participants were instructed to press a right click button asking for a micro-break, whenever through the task they felt fatigued equivalent to KSS≥5. When a micro-break was triggered by a right click, the task execution was interrupted after the earliest upcoming KSS pause (Fig 1). In the automatic sessions, the biofeedback system triggered the micro-break based on its prediction of KSS (explained further in the subsequent section) being ≥5 [42,43]. In this study, the micro-break consisted of a 25-s interruption of the task, while the participant took an active pause. During the micro-break, a down counter of the seconds from 25 to zero was displayed on the computer screen (Fig 1a). The green color has been shown to have restorative effects on attention and cognition [44,45]. The micro-breaks involved four repetitions of seated bilateral shoulder rotations with an elastic band where the shoulders were abducted horizontally up to 45°while keeping the elbows fixed around 90°. During the micro-break, the participants were also instructed to perform mindful breathing [46]. Besides the benefits of active pauses [47] especially during computer work [48], mindful breathing is associated with oxygenation and reduced mental load and stress to counteract sustained attention [46,49]. The breathing rate was at participants discretion, due to the diversity and individuality in breathing patterns [50].

### Familiarization and task engagement

The participants were instructed to perform the computer task and micro-breaks in four days prior to the first session. In addition, anthropometric measures, visual acuity, and general health and fatigue questionnaires were collected. Afterwards, the participants performed the computer task for 10-min. The participants received a brief overview of the experimental procedure also in the beginning of both sessions and performed the computer task for 5-min as an additional training before commencing the experimental protocol to reduce the learning effect. The participants were not informed about the principle of functioning of the biofeedback system. It was further explained that their choices of KSS do not affect their performance or automatic micro-breaks. To evaluate the perceived workload from the tasks, the questionnaire of National Aeronautics and Space Administration Task Load Index (NASA-TLX) [51] was administered after the task termination. The participants were informed that their performance was measured and compared with other participants to maintain motivation, and that achieving high performance make them candidates to win a monetary reward (100 Danish Kroner).

### The development of a statistical model for fatigue detection

To implement the biofeedback system, a statistical model of fatigue was developed based on previously collected oculometrics dataset (OLDSET) during an identical computer task [18]. The OLDSET consisted of the oculometrics extracted from gaze positions and pupil dilation, and KSS ratings from 38 participants in 40-min samples of an identical computer task without micro-breaks as described in [52]. The fatigue states for each segment were assigned based on KSS scores obtained after each segment. KSS scores were dichotomized a threshold of five, as KSS=5 corresponding to the statement of being “neither alert nor sleepy”, semantically implying the transition point between alertness and fatigue. Thus, the segments with the KSS value of ≥5 were assigned to a fatigued class and the KSS scores of <5 were assigned to an alert class. This dichotomization criterion has been used in previous studies, e.g. [42]. It is suggested as a critical value in the association between ocular metrics and sleepiness [43]. With this dichotomization criterion, 45% collected segments (205 out of 456) across the entire subject pool in the OLDSET turned out to be labeled as fatigued state.

Thirty-four features including oculometrics, sex, and age were used in setting up the classifier (Table 1). A series of viable classification models were examined as outlined in Table 2. A feature subset consisting of five features (denoted by asterisk in Table 1) was chosen using sequential floating forward feature selection [53]. The feature selection helps to choose a combination of features that best explain the separability of the two classes [53]. The classification criterion was the Youden’s J statistic or Youden’s index 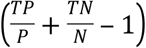 [54], where *TP, TN* are respectively the number of true positive (fatigued) and true negative (alert) instances, and *P* and *N* are the number of instances with respectively positive (fatigued) and negative (alert) labels. Here, 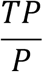 and 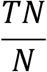 are respectively True Positive Rate (TPR), and True Negative Rate (TNR). The Youden’s index was computed using leave-one-person-out (LOPO) approach on a Random Forest model (Table 2) [55]. Different classifiers as outlined in Table 2 were examined using the selected feature subset as input and the class labels of fatigued or alert state as output. In LOPO approach, the classifier was trained using the data from all the participants except one, and it was tested using the excluded participant. This approach was performed for all the 38 participants to compute the average of classification performance across the entire participant pool. Finally, the ensemble of Decision Trees (DT Ensemble) was chosen based on its superior classification performance based on accuracy *ACC* = (*TP* + *TN*)/(*P* + *N*) (66±21%), TPR (61±29%), and TNR (70±22%) in comparison with the classifiers listed in Table 3, i.e. linear discriminant analysis, decision tree, k-nearest neighbors, support vector machines, Naïve Bayes, feed-forward neural networks, subtractive clustering-based Fuzzy classifier, Fuzzy c-means classifier, and Random Forest.

**Table 1.** The feature set.

| Feature | Description | Feature | Description |
| --- | --- | --- | --- |
| ***Blink-related Oculometrics*** |  | ***Fixation-related Oculometrics*** |  |
| BF* [Hz] | Blink Frequency | FF<sub>disp/dist</sub> [a.u.] | The ratio of the displacement to the distance between two successive fixations |
| BD [s] | Blink Duration | FD [s] | Fixation Duration |
| BGF [Hz] | The Frequency of Blinks accompanied by Gaze shifts [56] | FF [Hz] | Fixation Frequency |
| IBI [s] | Inter-Blink Interval (excluding IBI>20 s) | FF<sub>disp</sub> [cm] | Displacement between two successive fixations |
| LBF [Hz] | The frequency of long blinks (>200 ms) [57] | FF<sub>dist</sub> [cm] | Euclidean distance between two successive fixation centers |
| LBR [a.u.] | The ratio of long blinks (>200 ms) [57] to all blinks | OD [cm] | Overall Dispersion; the averaged Euclidean distance between fixation centers and center of fixations |
| DBF [Hz] | Double Blink Frequency (excluding IBI>700 ms) [58,59] | LFR [%] | The Rate of Long Fixation (>0.9 s) [60] to all fixations |
| BGR [a.u.] | The ratio of blinks accompanied by gaze shifts to all blinks [56] | ***Saccade-related oculometrics*** |  |
| TBS [s] | Time interval of <700 ms between a blink and its successive saccade | SVA* [s<sup>-1</sup>] | Saccade Peak Velocity Amplitude Relationship |
| PERCLOS* [%] | Percentage of the duration of closed eyes to opened eyes | SCD [s] | Saccade Duration |
| ***Pupil-related oculometrics*** |  | SF* [Hz] | Saccade Frequency |
| PD [mm] | Pupil Diameter | SPV [°/s] | Saccade Peak Velocity |
| PDIR* [mm] | Pupil Diameter Interquartile Range | SDA [s/°] | Saccade Duration Amplitude Relationship |
| PCV [a.u.] | Coefficient of Variation of Pupil diameter | SCR [°] | Saccade Curvature [39] |
| PH [a.u.] | Instantaneous phase of the pupil dynamics [61] | SA [°] | Saccade Amplitude |
| ***Demographics*** |  | SPD [°/s<sup>2</sup>] | Saccade Peak Deceleration |
| Age (continuous scale) |  | SPA [°/s<sup>2</sup>] | Saccade Peak Acceleration |
| Sex |  | ISI [s] | Short Inter-Saccade Intervals in <250 ms [62] |
|  |  | KPA [a.u.] | Kappa Coefficient of Ambient/Focal attention [63] |

\* The selected features for the classification model

**Table 2.** The details of the classification models

| **Model** | **Description** | **Configuration** | **Further considerations** |
| --- | --- | --- | --- |
| **LDA** | Linear discriminant analysis | Discriminant type: pseudolinear | Suitable to encounter singularity problems due to missing values [64] |
| **DT** | Decision Tree | Max number of splits: 7, Maximum number of categories: 10, Min parent size: 10, Prediction selection criteria: Curvature test, Pruning criterion: error, Min leaf size: 1, Split criterion: Gini's diversity index (GDI) | Choices for hyperparameters were based on recommended constraints [65] |
| **KNN** | k-Nearest Neighbors | 11-nearest neighbors classifier using the Euclidean distance metric and an exhaustive searcher | k=11 was chosen based on the highest classification performance in a grid search of k in [1 21], where the upper limit came from $\sqrt{n}$, where $n$ is the number of the training samples [66], Standardized feature set [67] |
| **SVM** | Support Vector Machines | Kernel: Gaussian | Standardized feature set [68] |
| **NB** | Naive Bayes | Predictor Distribution: Normal | Different kernels (Box, Traingular, and Epanechnikov) were tested. |
| **NN** | Feedforward Neural Network | One hidden layer including five neurons in the hidden layer with Scaled conjugate gradient backpropagation function. | Different number in of neurons from [1 10] was examined. 50 different random initial conditions and shuffled sequence of data presentation were applied. Standardized feature set [69] |
| **FIS-SC** | Fuzzy Inference System structure using subtractive clustering | cluster center's range of influence: 0.6, Input membership function: Gaussian, Output membership function: Linear | Cluster center's range was chosen in grid search between [0 1] with the steps of 0.1. Different membership functions have been tested |
| **FIS-FCM** | Fuzzy Inference System structure using Fuzzy C-Means clustering | FIS type: Sugeno, number of clusters: 2, input membership function: Gaussian, Output membership function: Linear | Different membership functions have been tested. [70] |
| **Fusion** | Major voting scheme | The same parameter setting for the LDA, DT, KNN, SVM, NB, NN, FIS-SC and FIS-FCM models used individually. | The single classifiers were combined as recommended in [71] |
| **DT Ensemble** | Ensemble of Decision Tree classifiers | Max number of splits: 5, Ensemble method: RobustBoost, Number of learning cycles: 50, Robust Error Goal: 0.25, Robust Max Margin: 1 | Robustness against noisy samples [72] due to subjectivity of class labels |
| **Random Forest** | Ensemble of Decision Tree classifiers | Max number of splits: 5, Number of trees:51 | Suitable for included categorical variables (sex) is included |

**Table 3.** The performance of the models to classify the fatigued state (Positive) from the alert state (Negative)

| **Model** | **Sensitivity** | **Specificity** | **Accuracy** |
| --- | --- | --- | --- |

|  | (TPR%) | (TNR%) | (ACC%) |
| --- | --- | --- | --- |
| **DT Ensemble** | 61 | 70 | 66 |
| **SVM** | 57 | 71 | 65 |
| **NB** | 61 | 68 | 65 |
| **Fusion** | 57 | 69 | 63 |
| **Random Forest** | 54 | 68 | 62 |
| **LDA** | 54 | 67 | 61 |
| **NN** | 53 | 67 | 61 |
| **FIS-FCM** | 54 | 67 | 61 |
| **FIS-SC** | 61 | 56 | 59 |
| **KNN** | 57 | 60 | 59 |
| **DT** | 56 | 52 | 54 |

The classification model with the best performance (i.e. DT-Ensemble) was picked to form the core of the biofeedback system. The DT-Ensemble with the configuration outlined in Table 2 was trained with the whole dataset consisting of 456 samples (38 participants × 12 segments) to make a statistical model to predict the class label of each segment of the biofeedback system.

The permutation test [73] was conducted on the OLDSET to further examine the classification accuracy of the DT Ensemble against the chance level accuracy obtained from 100 randomly permuted class labels. The sensitivity to the dichotomization criterion for the KSS was also performed on the OLDSET. The classification performance of the DT ensemble was significantly higher than chance level as assessed by the permutation test for the OLDSET (α=0.01). In addition, changing the dichotomization criterion of KSS scores to ≥6 as fatigued class for the DT Ensemble model did not lead to better classification performances than the criterion of five in the OLDSET.

During the computer task in the current study, the feature set was obtained across 20 consecutive cycles within a segment and the core of the biofeedback system classified the segment into either the fatigued or the alert class. The section titled “Oculometrics” outlines the performed analysis to obtain the oculometrics as features. If the segment was classified into the fatigued class, the biofeedback system triggered the micro-break command following the KSS pause after that specific segment. If the segment was classified into the alert class, no feedback was given. All aspects of the biofeedback system were implemented in MATLAB R2018a (The Mathworks, Natick, MA).

### Data acquisition and processing

A video-based monocular eye-tracker (Eye-Trac 7, Applied Science Laboratories, Bedford, MA, USA) coupled with a head tracker (Visualeyez II system set up with two VZ4000 trackers, Phoenix Technologies Inc., Canada) was utilized to measure the eye movements, pupil diameter, and point of gaze at a sampling frequency of 360 Hz. The coupling of the eye-tracker and the head-tracker was done using built-in software to integrate eye and head positioning data and to compensate for head movements allowing free head movements during the experiment. As reported by the manufacturer, spatial precision of the eye-tracker is lower than 0.5°of visual angle. The spatial accuracy is less than 2°in the periphery of the visual field. The calibration of the eye-tracker was performed before starting the task with 9-point calibration protocol and examined before the task execution and after the task termination on the calibration points. The measured accuracy was on average 0.7±0.4° across participants and did not significantly change across time (*p*>0.6). The experiments were conducted in a noise-and illumination-and temperature-controlled indoor room to rule out environmental confounding factors.

#### Oculometrics

Among all the features outlined in Table 1, the oculometrics were extracted from each segment. Saccades, blinks, and fixations were first identified for each segment using the algorithm of [74] as applied in [52]. Briefly, the algorithm initiated by the computation of visual angle between consecutive samples of point-of-gaze, followed by the its derivatives to the angular velocity and acceleration using a 19-samples-length second-order Savitsky-Golay filter [74]. It applied data-driven thresholds on the angular velocity to detect saccadic samples. Zero-valued samples of pupil diameter, corresponding to closed eyes or missing pupil image provided by the built-in software of the eye-tracker, constituted blink samples, and the rest of the samples were assigned to fixations. Pupil diameter (including linearly interpolated zero-valued samples) were filtered using a zero-phase low-pass third-order Butterworth to remove noise and artefacts [75]. Additional constraints were imposed to exclude invalid ocular events [52]. The data during the micro-breaks and KSS pauses were not included in the computation of oculometrics.

The frequency of blinks (BF), saccades (SF), and fixations (FF) were computed respectively as the number of blinks, saccades, and fixations during each segment divided by the duration of the segment. The mean duration of blinks (BD), fixations (FD), and saccades (SCD) were computed across each segment. Pupillary responses were characterized using the mean, coefficient of variation, interquartile range, instantaneous phase [61] of pupil diameter, respectively indicated by PD, PCV, PDIR, and PH. The number of closed-eyes samples (zero-valued pupil diameter) to opened-eyes samples was computed as PERCLOS. Blinks were further characterized by the frequency of blinks coincided by gaze shifts >2°(BGF) [56,76] and their ratio to the number of all blinks (BGR). The mean of inter-blink interval (IBI), the frequency of blinks occurring with IBI<700 ms (DBF), the number of long blinks >200 ms [57] to the segment duration (LBF), and the ratio of long blinks to all blinks (LBR). Saccades were further quantified in terms of the mean value of their peak velocity (SPV), amplitude (SA), curvature [39] (SCR), peak amplitude of saccadic acceleration (SPA) and deceleration (SPD) profiles, inter-saccadic intervals (ISI) excluding ISI>250 ms, and the slope of the line regressing peak velocity of saccades to their amplitude (SVA) and duration (SDA). Similarly, fixations were further characterized as the ratio of long fixations (>0.9 s) [60] to all fixations (LFR).

Gaze dispersion was characterized using the mean value of gaze-point displacements and distances between two successive fixations respectively computed as Euclidean distance between the center of gaze points of two successive fixations (FF_disp_), the summation of Euclidean distances between successive gaze-points from the onset to offset of saccades connecting the two successive fixations (FF_dist_), and the ratio of the FF_disp_ to FF_dist_ for the same two successive fixations (FF_disp/dist_). The successive fixations exceeding over feasible saccade duration of >100 ms in this study were excluded [77]. In addition, overall dispersion (OD) was quantified as the averaged Euclidean distance between fixation centers and center of fixations. The center of fixations, gaze points and fixation centers were obtained using the mean value of their corresponding coordinates. The dynamics of visual perception were also quantified using the kappa coefficient of ambient and focal attention (KPA) as defined in [63]. The mean of time intervals (<700 ms) between a blink and its successive saccade was also extracted as a feature in association with blink perturbation effects on saccades [78].

The selected features of SF, SVA, BF, PDIR and PERCLOS were computed in the biofeedback system in the same way as they computed from the OLDSET. To inspect the effect of the biofeedback system, the mean of the overall performance (OP) [52] across segments was computed. It represents how accurate and fast the pattern replication was done. Theoretically, the OP is a positive value with zero for the lowest performance.

### Statistical analysis

The statistical analysis was performed in SPSS 25. The classification performance of the deployed model (DT Ensemble) in the biofeedback system was reported in terms of the ACC, Sensitivity (TPR), and Specificity (TNR). The classification performance (ACC) was compared between the manual and automatic sessions using repeated measures analysis of variance (RM-ANOVA) and the interaction effect of the time of the day (morning or afternoon) on ACC was also considered. RM-ANOVA was also performed on the outcome variables (OP, KSS and oculometrics) with TOT (nine segments) and the automatic and the manual modes as within-subject factors (significance level *p*=0.05). Post-hoc comparisons between the segments were included in pairs indicated by Bonferroni correction. The Huynh-Feldt correction was applied if the assumption of sphericity was not met. The measure of effect size, partial eta-squared, 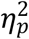, was also reported. The perceived workload (NASA-TLX scores) and the number of micro-breaks was compared between the automatic and manual sessions using paired t-test with the effect size in terms of Cohen’s *d* [79], to further evaluate the effectiveness of the biofeedback system. The normality of the variables was assessed using Shapiro–Wilk test. The KSS scores were transformed to normal distribution using square root transformation. The learning effect was analysed using RM-ANOVA to compare the OP across the first and second sessions.

## Results

In the automatic sessions, the model (DT Ensemble) identified the fatigued state from alert state with the following classification performance (Mean ± SD): ACC (69±16%), Sensitivity (59±35%), and Specificity (74±22%). The segments with the label of fatigued state constituted 55 segments in total of 180 segments. Fig 2 demonstrated the classification performance (ACC) of the model in both sessions for each participant. There was neither a significant difference in the classification performance between the automatic and manual sessions, nor an interaction of the time of the day on the ACC.

**Fig 2.**
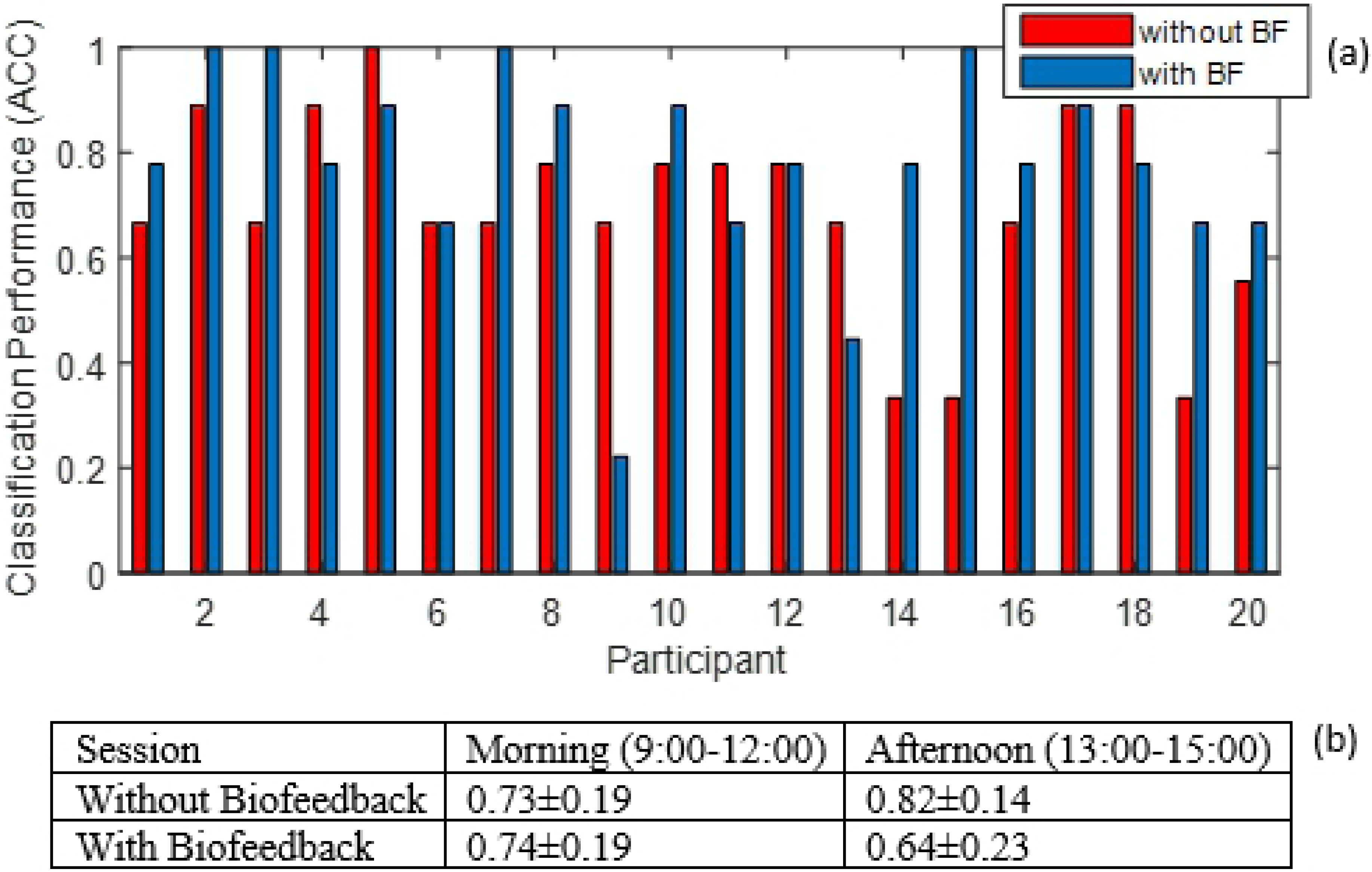
(a) Classification performance (ACC) of the DT Ensemble model for each participant in the manual and automatic sessions, (b) The ACC, Mean ± SD, in different time of the day (Morning and Afternoon)

The OP in the presence of the biofeedback did not significantly change, *F*(1,18) = 1.3, *p* = .262, 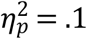, (Fig 3a). The OP increased significantly as TOT increased, *F*(8,152) = 4.7, *p* < .001, 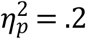. Pairwise comparisons revealed that the OP was significantly higher in segments eight and nine compared with two, five, and six, Fig 3a. In addition, there was no learning effect on the OP across the first and second sessions (Fig 3b).

**Fig 3.**
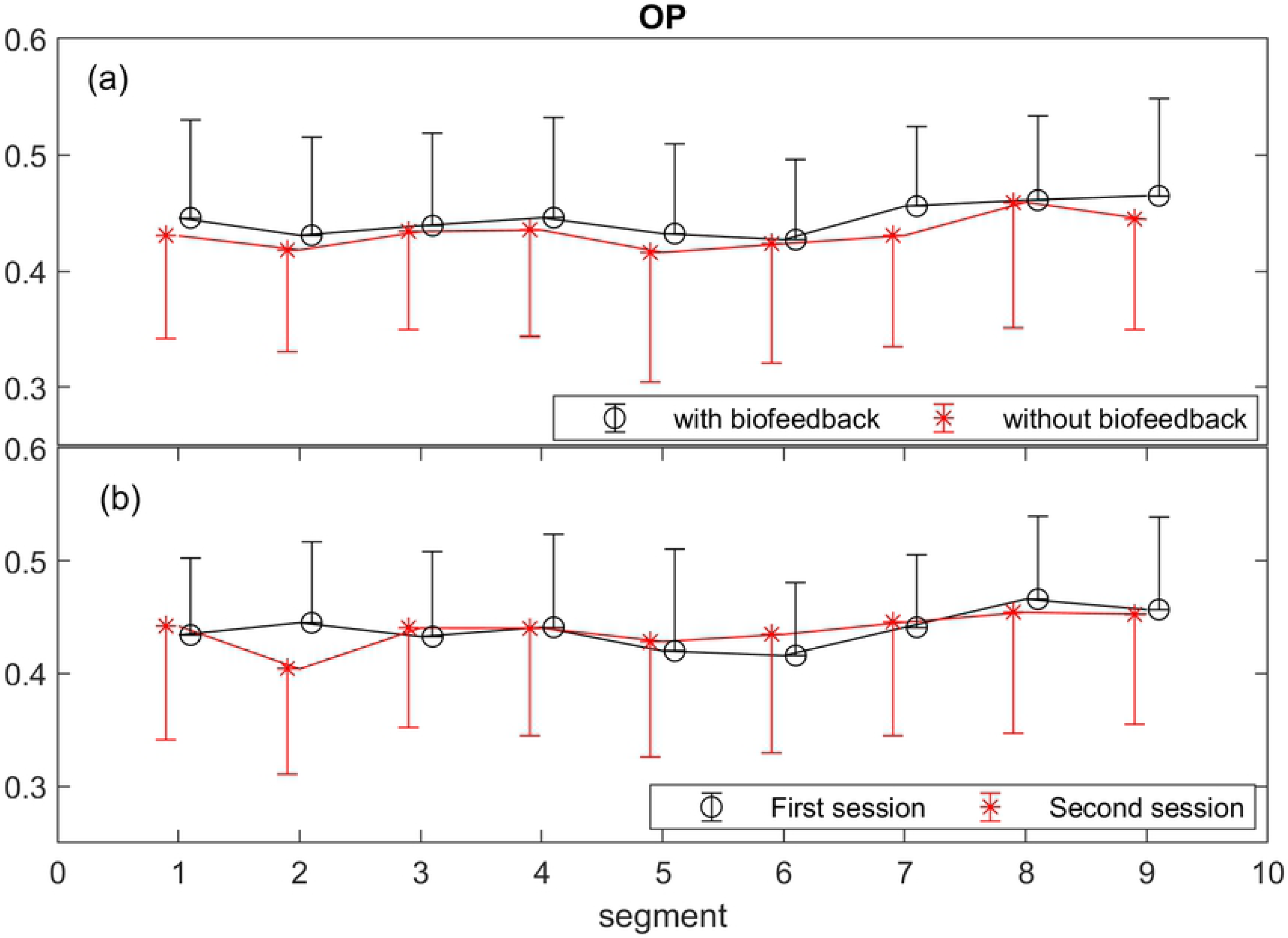
Comparison of the Overall Performance (OP) across sessions with and without feedback to assess biofeedback effect (a), and across the first and second sessions to inspect learning effect (b). The points and error bars respectively represent the mean and standard deviation values across the participants for each segment.

The participants reported significantly lower workload in the automatic (55±11) than the manual (65±8) sessions in terms of the total NASA-TLX scores, *t*(19) = 3.86, *p*=0.01, with the Cohen’s *d*=0.89 corresponding to a large effect size according to [80]. This improvement was more pronounced in mental and temporal subscales than the other workload subscales as demonstrated in Fig 4. There was no significant difference between the number of micro-breaks in the automatic sessions (2.9±1.9) from the number of micro-breaks in the manual sessions (2.5±2.3), *p*=0.55.

**Fig 4.**
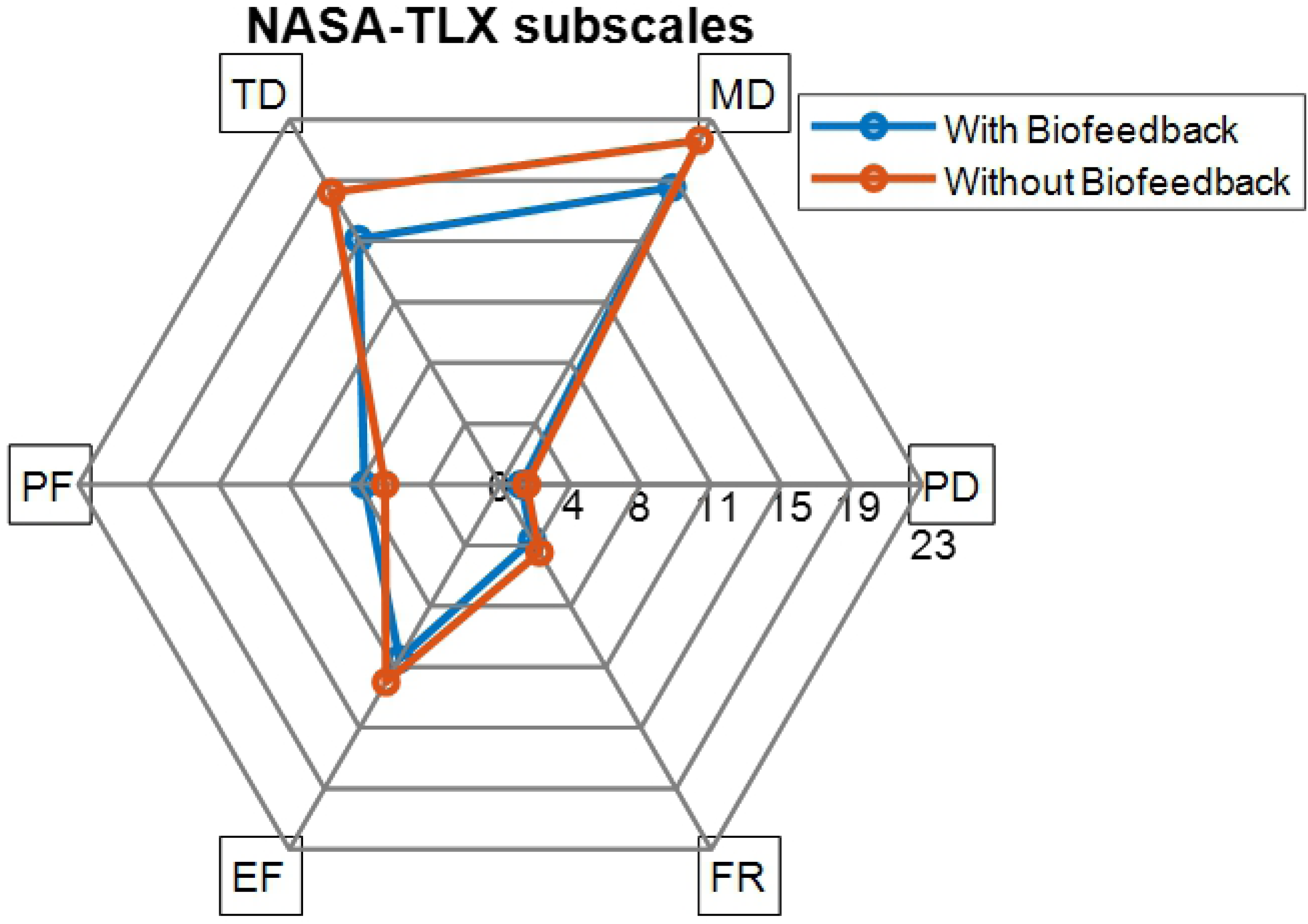
NASA-TLX scores in the task with biofeedback and without biofeedback

The KSS ratings significantly increased in both tasks with and without usage of biofeedback system as the segments increased *F*(5.8,109.6) = 15.6, *p* < .001, 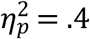, Fig 5. No significant change in the KSS scores was found between the automatic and manual sessions. However, a tendency of biofeedback×TOT interaction was found, *F*(5.9,113.6) = 1.7, *p* = .129, 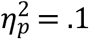. Pairwise comparisons showed that in the manual sessions, the KSS was lower in the first segment than in the segments 5-9, similarly between the segments 2-3 and 7-9, but in the automatic sessions, the significant difference was between the segments 1 and 2 being lower than both segments of 8 and 9.

**Fig 5.**
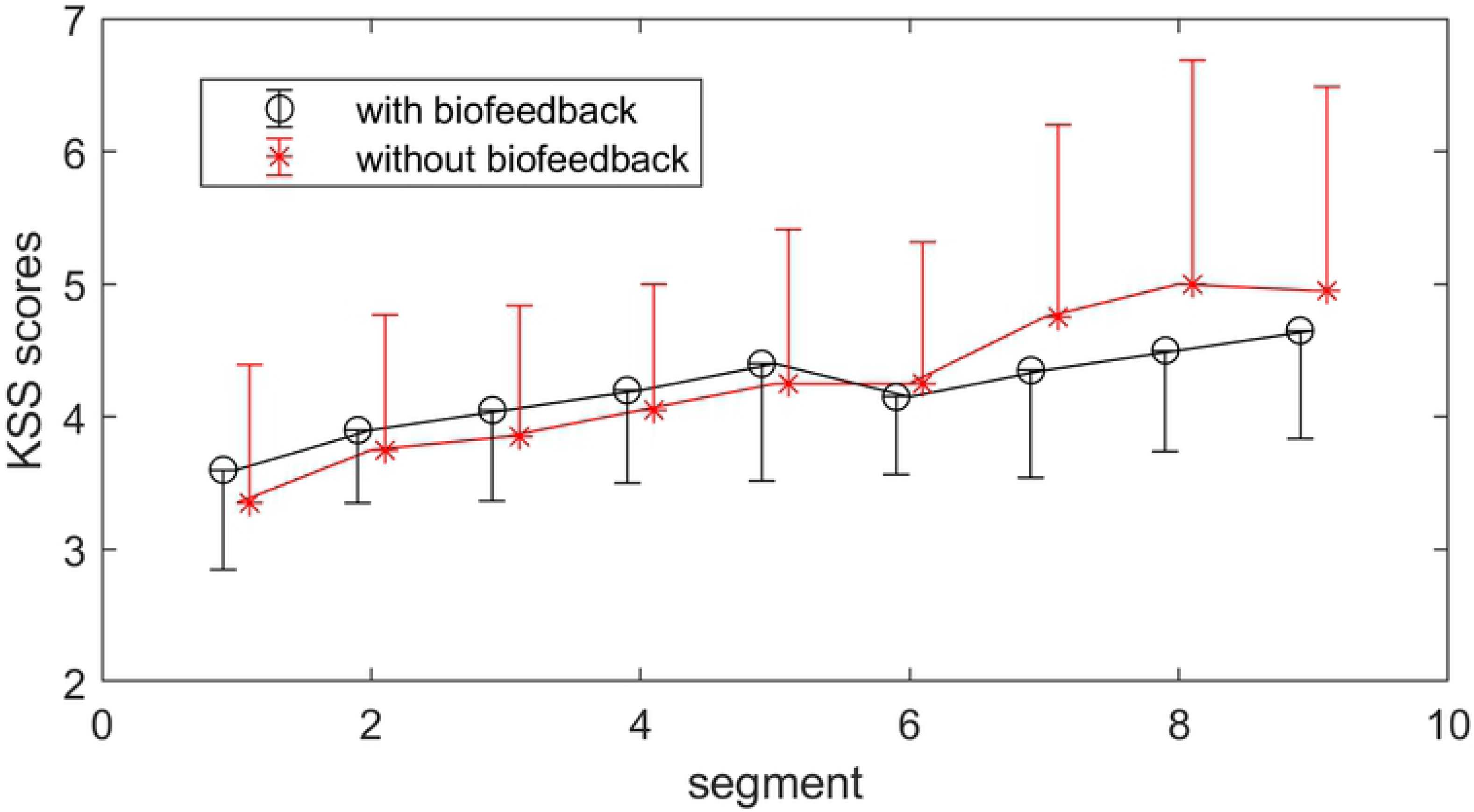
Subjective ratings of fatigue (KSS scores) in the tasks with and without using the biofeedback system. The points and error bars respectively represent the mean and standard deviation values across the participants for each segment.

Fig 6 shows the changes in the recruited oculometrics in the model to predict fatigue. The BF tended to increase with TOT, *F*(5.0,95.1) = 2.2, *p* = .058, 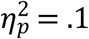. The PERCLOS increased significantly as TOT increased, *F*(8,152) = 2.3, *p* = .022,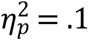. The SF decreased significantly as TOT increased, *F*(4.9,94.6) = 3.4, *p* = .007,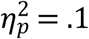. Pairwise comparisons revealed that the SF decreased significantly from segment 5 to 9. The SVA fluctuated significantly through TOT, *F*(8,152) = 2.2, *p* = .027,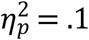. The change between segments 1 and 2 was significant in SVA. No significant effect of TOT on PDIR was observed. Neither any significant effect of biofeedback nor biofeedback×TOT interaction was found in any of the oculometrics.

**Fig 6.**
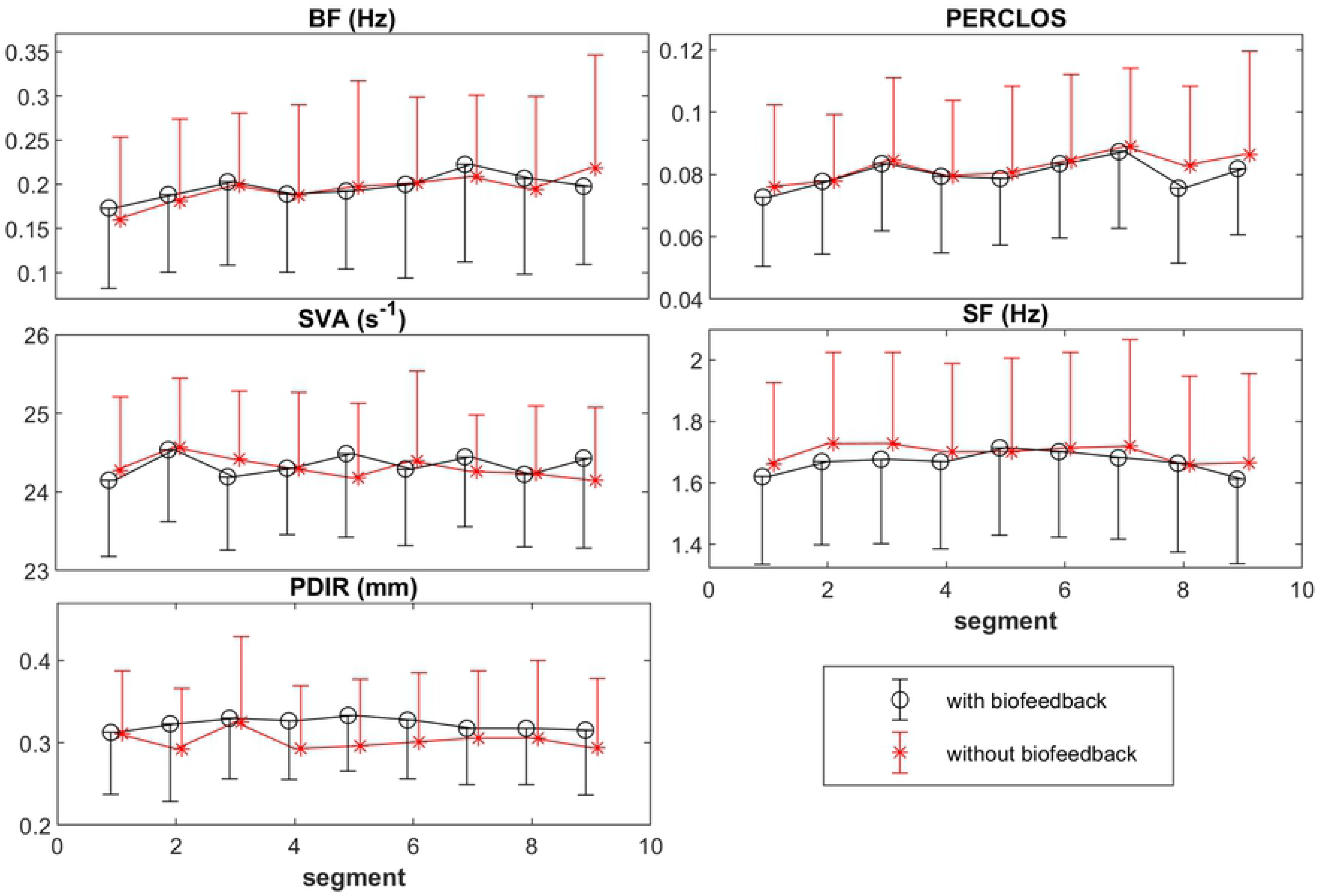
The changes through TOT in oculometrics recruited in the fatigue estimation model in the automatic sessions and manual sessions. The points and error bars respectively represent the mean and standard deviation values across the participants for each segment.

## Discussion

This study provided a novel framework to investigate the application of a biofeedback system reducing fatigue development in its early stages during computer work. The proposed biofeedback system deployed a statistical model of fatigue, which used quantitative features extracted from eye movements and pupillary responses, i.e. SF, PERCLOS, PDIR, BF, and SVA. The accuracy of the statistical model was promising considering the subjectivity of KSS scores. As hypothesized, the biofeedback system with the embedded micro-breaks, effectively counteracted fatigue development reflected in delayed trending towards fatigue (Fig 5) and decreased perceived workload (Fig 4).

The involved oculometrics (SF, PERCLOS, PDIR, BF, and SVA) have been previously reported to be reliable and sensitive to fatigue progression as well as mental load [18,39,81–84]. The PERCLOS and BF are reported to increase with fatigue [81,85], which is in line with the current results. The decrease in SF and increase in BF along-side with TOT were also in agreement with previous findings [86]. Saccadic main sequence and the range of pupil diameter decreases and increased, respectively with TOT [18,87], but the SVA and PDIR did not change monotonically with TOT, most likely because of the presence of micro-breaks.

To the best of our knowledge, this was the first study to deploy a statistical model of fatigue in a biofeedback system to trigger objective micro-breaks, and thereby to elaborate self-awareness of fatigue. A few studies have contributed in noninvasive fatigue detection. In [88], mental fatigue was detected offline using 31 statistical features from saccades, fixations, blinks, and pupillary responses exhibiting 77.1% accuracy with 10-fold cross validation via an SVM classifier. Our biofeedback system reached approx. 70% of accuracy. However, the accuracy of our method was achieved by the LOPO cross validation and used only five features (versus 31 in [81]), which may facilitate real-time applications. Nine 30-s data samples collected from each participant before and after two 17-min cognitive tasks, were used to detect mental fatigue in [88]. A numerical rating scale has been used as a subjective rating of fatigue in [88], however, the samples recorded before and after the cognitive tasks have been respectively labeled as non-fatigued and fatigued, regardless of individual differences in fatigue perception as opposed to the current study. A recent study using wearable electroencephalography has classified fatigue from alertness using an SVM classifier, based on KSS threshold of five, with the accuracy of about 65% in a 10-fold cross validation [89]. In [90], fatigue, subjectively labeled using a different rating scale, has been classified via a feedforward neural network using nine features extracted from computer user interactions with mouse and keyboard achieving an accuracy of 81% in a hold-out cross validation. The classification model proposed in [90], has been validated using the same group of individuals as opposed to the present study. Moreover, the features have been extracted over the period of one hour [90], which is much longer than a segment (≈200 s) to trigger micro-breaks questioning the practical use of such approach.

One general limitation in the study of fatigue is the inaccuracies of subjective ratings (KSS scores). Although it is still one of the most commonly used methods to acquire fatigue level [91], it could be affected by factors such as experimental design [92] and individual’s emotional state [93][94]. One may suggest the task performance (OP) as an alternative to KSS. However, OP cannot necessarily be translated into fatigue levels in early detection of fatigue, Fig 2a. Additionally, the task performance may consistently change with TOT [7].

Another important issue to consider is the effect of circadian rhythms on the accuracy of the fatigue state estimation [95]. Circadian rhythm is a source of variability in oculometrics [96,97] and cognition [98], which makes the prediction of fatigue quite challenging. The non-significant difference between the classification accuracy of the DT Ensemble model for the half of the participants who did the tasks in the morning (9:00-12:00) and the rest of the participants who did the tasks in the afternoon (13:00-15:00) confirmed the robustness of the model to psychophysiological changes in response to circadian rhythms, (cf. Fig 2b).

An efficient and effective design for micro-breaks is quite challenging especially due to the complex interferences between physical and mental demands of a task [99,100]. Interestingly, reduced perceived workload was observed in the sessions where the micro-breaks were triggered by the biofeedback system compared with the manual sessions. Slight improvements of task performance (OP), Fig 3a, and delayed inclination to fatigue, Fig 5, were observed through using the biofeedback system. Even though the improvement in the performance was statistically insignificant, one may conceive that in a long run the slight improvement in the performance may be of importance for the prevention of musculoskeletal disorders [14,15,101].

Micro-breaks were applied in this study based on important considerations. The activities during micro-breaks should not demand for the same mental resources that a task might require [102]. Considering the multiple resource model [103], targeting the same mental resources may decline performance. Accordingly, in comparison with the task demands, the micro-breaks intuitively required little physical and mental demands and perhaps a low vigilance to attend to down-counter displayed on the screen. In practice, to avoid too frequent and invalid micro-breaks, interactive micro-breaks [27] and model adaptation is suggested to study through the presented framework. In comparison a previous study of [104], the simplicity and effectiveness of the proposed micro-break as well as the unconstrained technique of eye-tracking potentially meet constraints of out-of-lab settings.

## Conclusion

In line with our hypothesis, this study shows for the first time that the integration of oculometrics-based biofeedback in the design of micro-breaks is robust against circadian variations and effective in fatigue mitigation during computer work. The effectiveness of the oculometrics-based biofeedback was evidenced by the decreased perception of workload and further by the delayed inclination to fatigued state using the biofeedback system compared with self-triggered micro-breaks. In sum, the use of oculometrics as objective indices of fatigue in a biofeedback system may be a viable approach to impede fatigue development.

